# Correlated motions of conserved polar motifs lay out a plausible mechanism of G protein-coupled receptor activation

**DOI:** 10.1101/2020.01.27.920769

**Authors:** Argha Mitra, Arijit Sarkar, Márton R. Szabó, Attila Borics

## Abstract

Recent advancements in the field of experimental structural biology have provided high-resolution structures of active and inactive state G protein-coupled receptors (GPCRs), a highly important pharmaceutical target family, but the process of transition between these states is poorly understood. According to the current theory, GPCRs exist in structurally distinct, dynamically interconverting functional states of which populations are shifted upon binding of ligands and intracellular signaling proteins. However, explanation of the activation mechanism on an entirely structural basis gets complicated when multiple activation pathways and active receptor states are considered. Our unbiased, atomistic molecular dynamics simulations of the mu-opioid receptor in a physiological environment revealed that external stimulus is propagated to the intracellular surface of the receptor through subtle, concerted movements of highly conserved polar amino acid side chains along the 7^th^ transmembrane helix. To amend the widely accepted theory we suggest that the initiation event of GPCR activation is the shift of macroscopic polarization between the ortho- and allosteric binding pockets and the intracellular G protein-binding interface.

Table of Contents Graphic

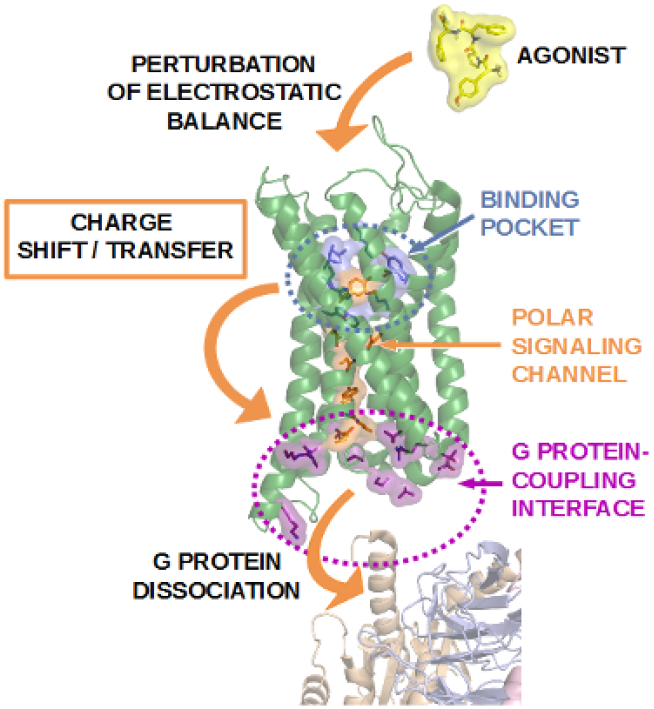

## INTRODUCTION

G protein-coupled receptors (GPCRs) are located on cell surfaces and act as communication interfaces for external stimuli exerted by structurally diverse molecules. Upon activation GPCRs initiate signal transduction through interactions with G proteins and arrestins, and control a variety of intracellular processes. Owing to this, approximately 34% of all prescription pharmaceuticals target members of this receptor family.^1^ However, application of such drugs is often limited by a number of unwanted side effects due to non-selective activation of multiple GPCRs, or multiple signaling pathways associated with one receptor. The most recent challenge of rational drug design is, therefore, to develop signaling pathway-specific, or in other words ‘functionally selective’ GPCR agonists. To address this challenge, complete understanding of the structural mechanism of GPCR activation is necessary. Opposed to the high diversity of external activators, signaling is mediated by only a few types of G proteins, advocating that GPCR activation may follow a general mechanism.

The structure of GPCRs consists of a conserved bundle of seven transmembrane α-helices and highly dynamic extracellular and cytosolic domains of various lengths. High-resolution experimental structures are available for many GPCRs both in the active and inactive states (see http://gpcrdb.org),^2^ but the mechanism of transition between these forms is intensely debated. The most conspicuous difference between the active and inactive class A GPCR structures, published to this date, is a notable disposition of the 6^th^ transmembrane helix (TM6).^3–4^ However, such large dispositions were shown to occur even in the absence of a bound ligand, due to the inherent dynamics of the receptor structure, or may originate from the applied conditions of crystallographic structure determination, namely the attachment of fusion proteins or the application of crystallization chaperones.^5–7^ Apart from TM6 disposition, a possible role of intracellular loop 1 (ICL1) and the cytosolic helix (H8) in the activation mechanism was highlighted by dynamic NMR measurements of the mu-opioid receptor.^8^ Comparison of the structures of active and inactive state mu- and delta-opioid receptors suggested that an extended network of polar amino acids and water molecules connects the orthosteric ligand binding pocket to the cytosolic domains which may be functionally relevant. The highly conserved E/DRY, NPxxY and CWxP functional motifs as well as allosteric Na^+^ binding sites have been specified to participate in the activation mechanism of class A GPCRs.^9^ Conceivably, activation signal is transmitted to the intracellular surface of the receptor through the interplay of these microswitches, however, no direct evidence of such integral mechanism has yet been given. Real-time observation of such processes using conventional experimental techniques is unattainable.

A significant part of the now widely accepted theory of GPCR activation was provided by landmark molecular dynamics (MD) simulation studies.^3–4,10–11^ According to this theory, GPCRs exist as a dynamic ensemble of multiple active, inactive and intermediate states. The populations of active states is increased by agonist binding and the stabilization of an active structure facilitates the insertion of G proteins.^10^ The growing number of evidence of pre-coupled GPCR-G protein complexes in the absence of ligands, however, presents a challenge to the above hypothesis.^12^ In general, explanations given on an entirely structural basis are often diffuse and fail to provide unequivocal suggestion for a possible structural mechanism of GPCR activation, especially when multiple active states, or structurally similar but functionally different ligands are considered. Further limitations of previous MD studies are that simulation systems were confined to the TM region of GPCRs, embedded in very simplistic representations of the cell membrane. Most recently, special effects of charged interfacial lipids on β_2_-adrenergic receptor signaling was demonstrated, drawing attention to the importance of accurate membrane representation.^13^

The mu-opioid receptor is one of the most extensively studied GPCRs therefore appropriately represents the general structural features of class A (rhodopsin-like) GPCRs. In order to get a deeper insight, we have performed all-atom MD simulations of the mu-opioid receptor on a μs timescale, in a physiologically relevant setup. Simulations of the active and inactive receptors were executed in caveolar membrane environment, in the presence of the endogenous agonist endomorphin-2 (EM2)^14^ and the G_i_ protein complex, or beta-arrestin-2. In addition, control simulations were carried out with fused T4-lysozyme or intacellularly bound Nb39 nanobody, representing the previously applied crystallization conditions.^5,7^

## METHODS

### System building

The crystallographic structures used in this study were downloaded from the Brookhaven Protein Data Bank (http://www.rcsb.org): active mu-opioid receptor (pdb code: 5C1M), inactive mu-opioid receptor (pdb code: 4DKL), heterotrimeric G_i_ protein complex (pdb code: 1GP2), Nb39 nanobody (pdb code: 5C1M), T4-lysozyme (pdb code: 4DKL), β_2_-adrenergic receptor complexed with G_s_ protein (pdb code: 3SN6) and rhodopsin complexed with beta-arrestin-2 (pdb code: 4ZWJ). These latter two structures were used as templates to orient the G_i_ protein complex and beta-arrestin-2 to the mu-opioid receptor. The full sequence of the murine mu-opioid receptor (UniProtKB-P42866-OPRM1) was obtained from UniProt (http://www.uniprot.org), and the coordinates of the membrane orientation from the OPM server (http://opm.phar.umich.edu) The crystallization chaperone and fusion protein (Nb39 nanobody and T4-lysozyme, respectively) were removed from the crystallographic structures. The Swiss-PdbViewer was used to retrieve all missing, modified or mutated residues of the transmembrane (TM) domain of the receptor. GTP was generated in CHARMM-GUI (http://www.charmm-gui.org) and edited manually to replace GDP in the G_i_ complex.

The exclusion of the drag posed by the mass of the N- and C-terminal domains may provide unrealistic results for the dynamics of transmembrane helices. To model the missing N- and C-terminal domains 10 ns folding simulations of the N- and C-terminal domains were performed using the GROMACS ver. 5.1.4 program package, the AMBER ff99SB-ILDN-NMR force field and GB/SA implicit solvation. During MD simulations, the system temperature was set to 310 K and maintained by the v-rescale algorithm. Ten parallel simulations were run for both the N-and C-terminal domains from where the resultant, folded structures were evaluated and selected based on the compactness, accessibility of post-translational modification- and TM region attachment sites. Glycosylation sites were predicted using the NetNGlyc 1.0 online server.^15^ The selected domain structures were linked to the TM region using Pymol ver. 2.1.0.

Four intracellular partners were used in this study, namely the heterotrimeric G_i_ protein, beta-arrestin-2, Nb39 nanobody and T4-lysozyme. Among them the last two were used as control reference simulation systems. The first three proteins were attached non-covalently to the receptor while T4-lysozyme was fused with the receptor replacing the third intracellular loop (ICL3), similarly to that in the crystallographic structure of the inactive mu-opioid receptor (pdb code: 4DKL).

CHARMM-GUI was used to include various post-translational modifications as well as to build membrane bilayers. Complex type glycans were added to the N-terminal domain, containing a common core (Manα1-3(Manα1-6)Manβ1-4GlcNAcβ1–4GlcNAcβ1–N) and sialic acid (N-acetylneuraminic acid) at glycosylation prone N9, N31 and N38 residues of the N-terminal domain.^16^ Phosphorylation of S363 and T370 were done for all the complexes, while S375, T376 and T379 sites at the C-terminal domain were phosphorylated in addition for the arrestin complexes.^17^ The C170 residue of ICL2 was palmitoylated.^18^

A caveolar membrane environment, considered to be the physiological environment of the mu-opioid receptor, was built using the membrane builder tool of CHARMM-GUI. CHARMM36 parameters were used to build complex, multicomponent membrane systems which included cholesterol (CHL-32.8%), 1-palmitoyl-2-oleoyl-glycero-3-phosphocholine (POPC-14.9%), 1-palmitoyl-2-oleoyl-sn-glycero-3-phosphoethanolamine (POPE-27.8%), 1-palmitoyl-2-oleoyl-sn-glycero-3-phospho-L-serine (POPS-3.6%), 1-palmitoyl-2-oleoyl-sn-glycero-3-phosphoinositol (POPI2-6%), palmitoyl-sphingomyelin (PSM-9.9%) and monosialodihexosylganglioside (GM3-5%).^19^ GM3 gangliosides were generated separately using the glycoprotein builder tool of CHARMM-GUI, and then added manually to the membrane. The asymmetric upper and the lower leaflet membrane compositions were specified in a most probable ratio.^20^ The membrane builder was also used to embed the glycosylated, palmitoylated and phosphorylated full sequence model of the mu-opioid receptor into the membrane. Systems were then solvated explicitly with TIP3P water molecules in a hexagonal shaped periodic box, and sodium and chloride ions (0.15 M) were added to neutralize the net charge and to attain physiological ionic strength. System coordinates and topologies were generated in GROMACS format.

EM2^14^, a putative peptide agonist of the mu-opioid receptor was built manually using Pymol ver. 2.1.0. The binding site was confirmed by flexible docking of this ligand to the active state mu-opioid receptor crystallographic structure (pdb code: 5C1M) using the Autodock ver. 4.2 software and the Lamarckian genetic algorithm. All ϕ, ψ, and χ^1^ ligand torsions, as well as receptor side chains in contact with the bound ligand ^5^ were kept flexible. Docking of EM2 was performed in an 8.0 nm × 8.0 nm × 8.0 nm grid volume, large enough to cover the whole binding pocket of the receptor region accessible from the extracellular side. The spacing of grid points was set at 0.0375 nm and 1000 dockings were done. The resultant ligand-receptor complexes were clustered and ranked according to the corresponding binding free energies. The lowest energy bound state was selected for simulations, in which specific ligand-receptor interactions observed in the crystallographic structures were present. Cryo-electron microscopic structure of the mu-opioid receptor and the peptide agonist DAMGO, published later, have confirmed the correct orientation of EM2.^6^

### MD simulations

All equilibration and production MD simulations were performed using the GROMACS ver. 5.1.4. molecular dynamics program package. Eight independent simulations were performed, four each for inactive and active mu-opioid receptors, complexed with heterotrimeric G_i_ protein, beta-arrestin-2, Nb39 nanobody and T4-lysozyme. After orienting and adding EM2, the resultant complex systems were energy minimized thoroughly performing 5000 steps steepest descent, followed by 5000 steps conjugate gradient minimization with convergence criteria of 1000 kJ/mol nm^−1^ in both cases. After minimization, systems were subjected to a six-step equilibration protocol of positionally restrained MD simulations in the NVT and then, after 2 steps, in the NPT ensemble at 303.15 K and 1 bar, having the positional restraints on the heavy atoms of the proteins and membrane constituents decreasing gradually. The first three equilibration MD runs were done for 25 ps in 1 fs time steps, and the following three was continued for 100 ps in 2 fs time steps. The following, further parameters were applied: the LINCS algorithm was used to constrain all bonds to their correct length, temperature was regulated by the v-rescale algorithm with a coupling constant of 1 ps, semi-isotropic pressure coupling was applied with a coupling constant of 5 ps and compressibility of 4.5 × 10-5 bar^−1^. The PME method was used to calculate energy contributions from non-bonded interactions with all cut-off values set to 1.2 nm. After equilibration, production simulations were performed for 1 μs at 310 K in the NPT ensemble, with other parameters same as above. The coordinates were stored in every 5000th steps yielding trajectories of 100.000 snapshots.

### MD trajectory analysis

Analysis of MD trajectories was performed using the analysis suite of the GROMACS 5.1.4 package. The analyses of MD trajectories were performed to evaluate protein conformational changes, stability of the molecular complexes, as well as to investigate previously described interactions and their role in different activation states of the receptor.

For each MD trajectory root mean square deviation (RMSD) calculations were carried out to assess the structural stability of the complex and demonstrate significant displacements of structural components as a function of time. RMSD values of protein backbone atoms were calculated in comparison with the active and inactive state starting structures. Conformational fluctuations of specific amino acid side chains were analyzed by measuring side chain χ^◻^ angles and calculating the frequency of transitions between rotameric states using gmx chi. The gmx helix utility was used to calculate helix properties. Secondary structure assignment was done using the DSSP method.^21^

The occurrence and frequency of intra- and intermolecular H-bonds were calculated using the gmx hbond utility. The donor-acceptor distance and donor-hydrogen-acceptor angle cut-offs for H-bond assignments were set to 0.35 nm and 60.0 degrees, respectively. The presence of salt bridges was monitored by measuring distance and angle between the corresponding acidic and basic side chain functional groups using gmx distance and gmx angle, respectively. The distance threshold for salt bridge assignment was 0.4 nm and the angle threshold was 90.0 degrees.

The extent of correlation of atomic displacements was examined by dynamic cross-correlation matrix analysis (DCCM) integrated into an earlier version of the GROMACS suite (g_correlation, ver. 3.3).^22^ The Gimp ver. 2.8 software was used for image analysis of the obtained DCCM maps, where the extent of correlation was demonstrated by color intensity. The threshold of assignment of correlation was red color intensity corresponding to >0.7 MI (mutual information). Amino acid side chains having at least 4 atoms participating in correlated motions were considered.

Systems were visualized using Pymol ver. 2.1.0 or VMD ver. 1.9.3 softwares and graphs were prepared using the Xmgrace ver. 5.1.25 program.

### Sequence alignment and conservation analysis

244 sequences of class A mouse GPCRs (without orphan and olfactory receptors) were retrieved from the UniProt database in FASTA format. The Clustal Omega program^23^ was used to align those multiple sequences and the results were analyzed using Jalview ver. 2.10.5. The OPRM_MOUSE (P42866) sequence was set as reference. The sequences were compared based on percentage of identity.

## RESULTS AND DISCUSSION

Atomic displacement analysis of transmembrane helical backbones indicated, that TM6 assumed intermediate conformations during simulations with minor changes from the corresponding starting structures (Figure S1). This is in line with previous simulation results, where notable TM helix rearrangements were only observed at longer timescales and in the absence of bound intracellular proteins.^10^ Initial conformations of ICL1 and H8 were maintained throughout simulations in each receptor state, regardless of the intracellular interacting partners (Figure S2-S3). The second intracellular loop (ICL2), on the other hand, adopted a stable α-helical structure when bound by beta-arrestin-2 and partially unfolded upon interaction with the G_i_α subunit, independent of the state of the receptor (Figure S4). In the active states, increased frequency of intermolecular hydrogen bonds was observed involving ICL2, helix 5 of G_i_α and the finger- and C-loops of beta-arrestin-2 (Figure 1, Table S1). These observations indicate that the conformations of ICL1 and H8 are controlled by the receptor state, whereas ICL2 adapts its structure to the bound signaling proteins, therefore may be partially responsible for signaling pathway specificity. Such dynamics of ICL2 was not indicated by the published high resolution structures of this receptor.^5–7^ Secondary structure analysis of the control systems with Nb39 or T4-lysozyme fusion suggests that these systems better represent the arrestin-bound state of the receptor (Figure S4).

**Figure 1.**
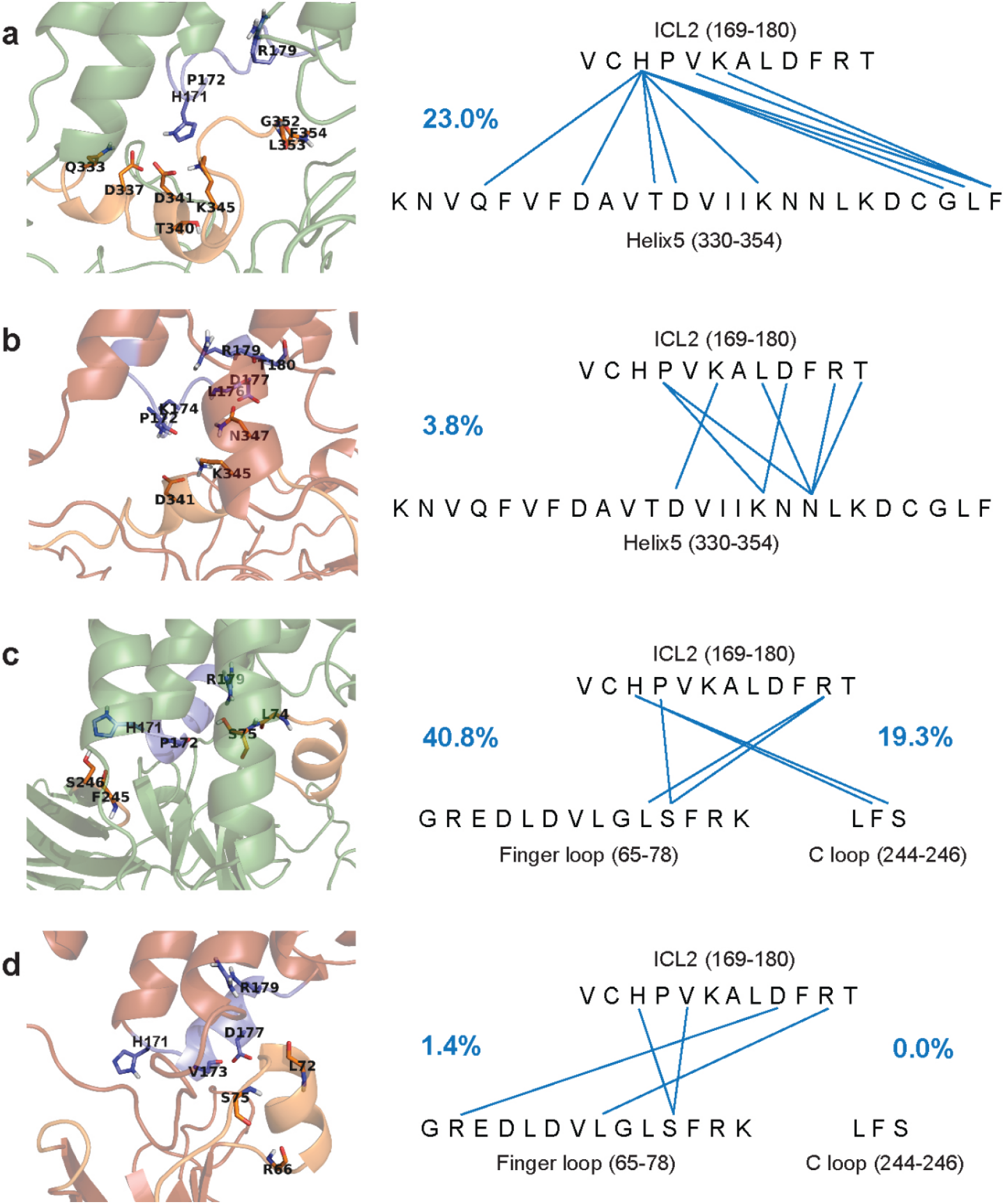
Frequency and donor and acceptor sites of intermolecular H-bonds between ICL2 of the mu-opioid receptor and the G_i_ protein or beta-arrestin-2. Frequency of H-bonds are expressed as percentages of the total structural ensemble and indicated by blue numbers. ICL2 is shown as blue cartoon and sticks. Helix 5 of the G_i_α subunit and the finger- and C loops of beta-arrestin-2 are shown as orange cartoon and sticks. (a) Active receptor and the G_i_α subunit. (b) Inactive receptor and the G_i_α subunit. (c) Active receptor and beta-arrestin-2. (d) Inactive receptor and beta-arrestin-2.

Analysis of intramolecular salt bridges and H-bonds between conserved motifs (Table 1) indicated, in agreement with previous proposals,^7^ that interactions between D164^3.49^ and R165^3.50^ of the DRY motif were more frequent in the inactive states. Specific role of the previously reported DRY-TM5,^7^ DRY-TM6,^7^ CWxP-TM7^24^ and NPxxY-TM^5^ contacts in the activation mechanism was not substantiated by the simulation results. No systematic connection was found between the frequency of those interactions and physiologically relevant receptor states and complexes. A salt bridge between R165^3.50^ (DRY) and D340^8.47^ (H8), however, was found to be present only in the active state and most frequent in the presence of the G_i_ protein complex. This specific interaction was not described previously as, opposed to the above mentioned contacts, it was not evidently present in the reported high resolution structures.^5–7^ Our data suggest that this contact may be important for receptor activation and it is further supported by earlier mutation experiments.^25^

**Table 1.**
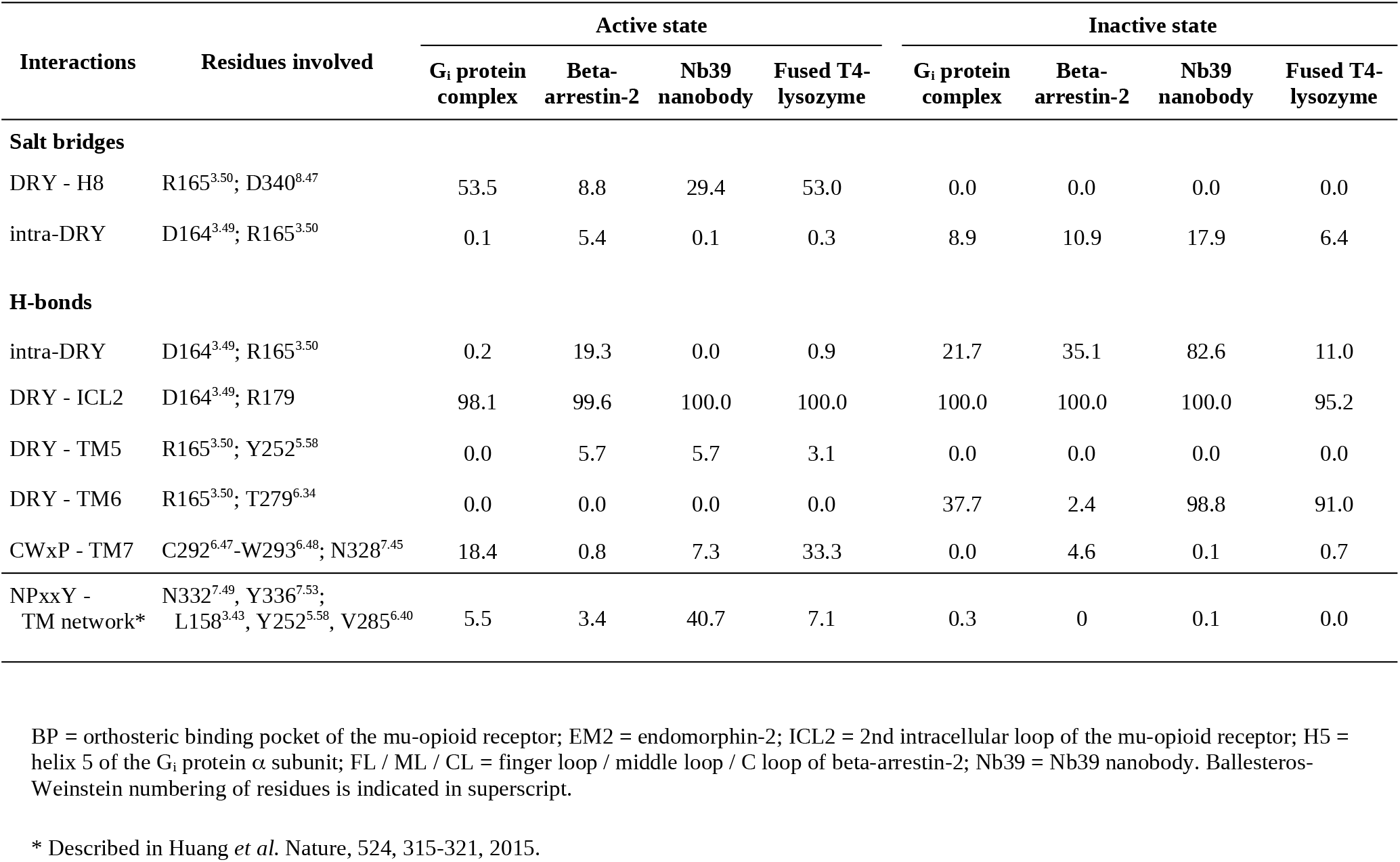
Frequency of intramolecular salt bridges and H-bonds expressed as percentages of the total conformational ensemble, generated by MD simulations.

Dynamic cross-correlation analysis of the transmembrane domain and the extra- and intracellular loops provided the most striking results (Figure 2). According to those, the orthosteric binding pocket is connected to the intracellular surface through a channel of polar amino acid residues, of which motions are highly correlated (Figure 3). Such concerted motions were observed only for the active receptor–G_i_ protein complex (Figure 3, Figures S5-S9), suggesting that this phenomenon has a fundamental role in G protein-mediated signaling. Residues of this polar signaling channel are located mostly on the 7^th^ transmembrane helix (TM7), in the inner region of the transmembrane helical bundle, most distant from the surrounding membrane environment. All channel residues are parts of highly conserved functional motifs and allosteric Na^+^ binding sites, except for Y326^7.43^ of the binding pocket and N340^8.47^ at the G protein-binding interface. The increased variability of these residues is associated with ligand and G protein specificity, respectively. Analysis of the individual dynamics of these specific side chains revealed that the observed movements are small and mostly occur without the transition between rotameric states (Table S2). Considering that the orientation of amino acid side chains in the orthosteric binding pocket of the mu-opioid receptor are nearly identical in the agonist-^5,6^ and antagonist-bound states^7^, our results suggest that the underlying event of receptor activation is the change of macroscopic polarization in a shielded central duct of the transmembrane domain. Although classical force field methods cannot provide quantitative details of such mechanism, independent mutation data provides direct evidence for the interplay of these polar and charged amino acid side chains during receptor activation. Impaired G protein signaling or elevation of constitutional activity was observed for mutant receptors, where residues of the above mentioned polar signaling channel were replaced,^9,24–32^ while receptor activity was preserved in double mutants, where the net charge of channel residues was kept intact.^28^

**Figure 2.**
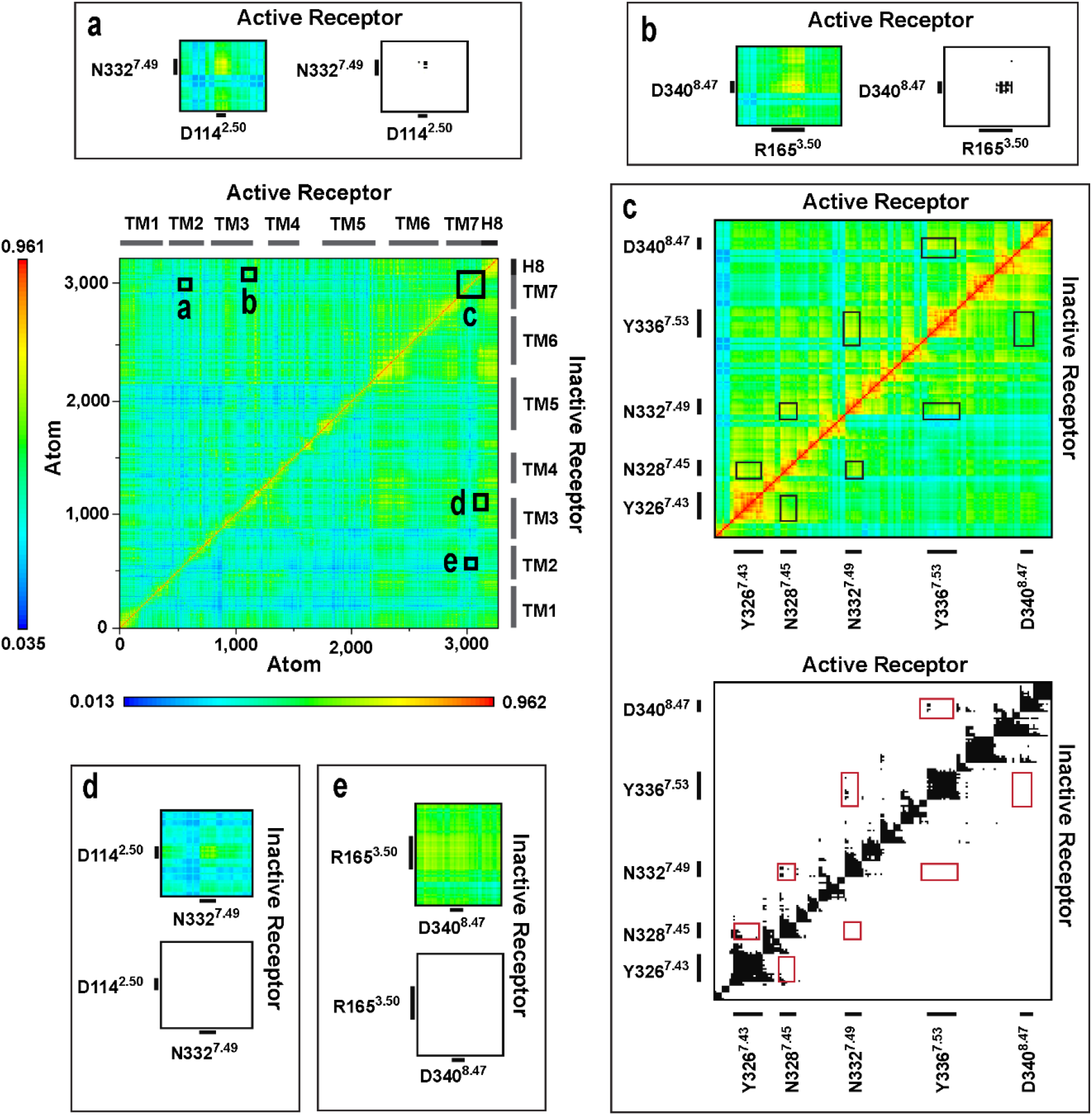
Dynamic cross-correlation matrices of the G_i_ protein-bound mu-opioid receptor in active and inactive states. Panels (a-e) are magnified views of regions of amino acid residues of interest. Black and white panels show correlations above the threshold of 0.7 MI.

**Figure 3.**
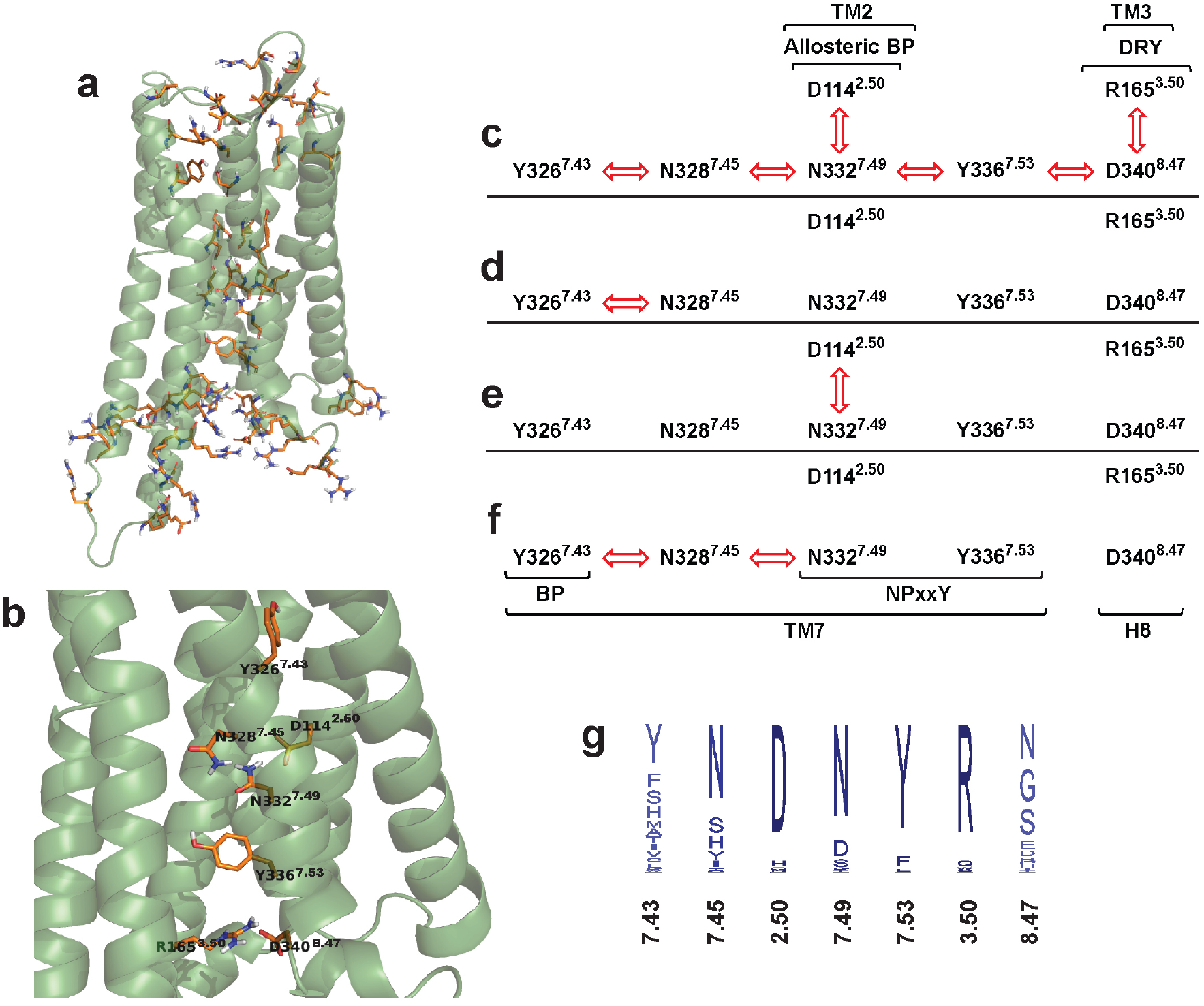
The polar signaling channel of the mu-opioid receptor revealed by dynamic cross correlation analysis. (a) Polar amino acids of which motions are correlated in the G_i_ protein-bound active state. (b) Polar amino acids of which motions are correlated and connecting the orthosteric binding pocket to the G protein-binding interface. Diagrams of channel residues in (c) the active receptor – G_i_ protein, (d) inactive receptor – G_i_ protein, (e) active receptor – beta-arrestin-2 and (f) inactive receptor – beta-arrestin-2 complexes. Red arrows indicate correlated motions of the respective amino acids. (g) Degree of conservation of polar signaling channel residues of class A GPCRs. Non-polar hydrogens are omitted for clarity.

The shift of macroscopic polarization may be assisted by the inherent dipole moments of TM helices. Generally, a more ordered α-helical segment possesses a higher dipole moment, which can participate in various conduction processes.^33^ Analysis of the evolution of helix properties during simulations exhibited that TM7 is the most ordered among the TM helices of the active, G_i_ protein-bound receptor. Furthermore, the helicity of TM7 is closest to ideal when the receptor is G_i_ protein-bound and least ideal when complexed by beta-arrestin-2 (Figure 4). This accentuates the role of TM7 in the activation mechanism and it is corroborated by previous reports.^9, 34^ The role of electrostatic forces and the importance of charge balance is further supported by the known effect of elevated concentrations of Na^+^ ions^35^ and the concept of voltage sensing.^36–37^ According to this latter, changes in the transmembrane electrostatic potential (V_m_) resulting from the rearrangement of charged species and polar membrane components could elicit functional effects in GPCRs.

**Figure 4.**
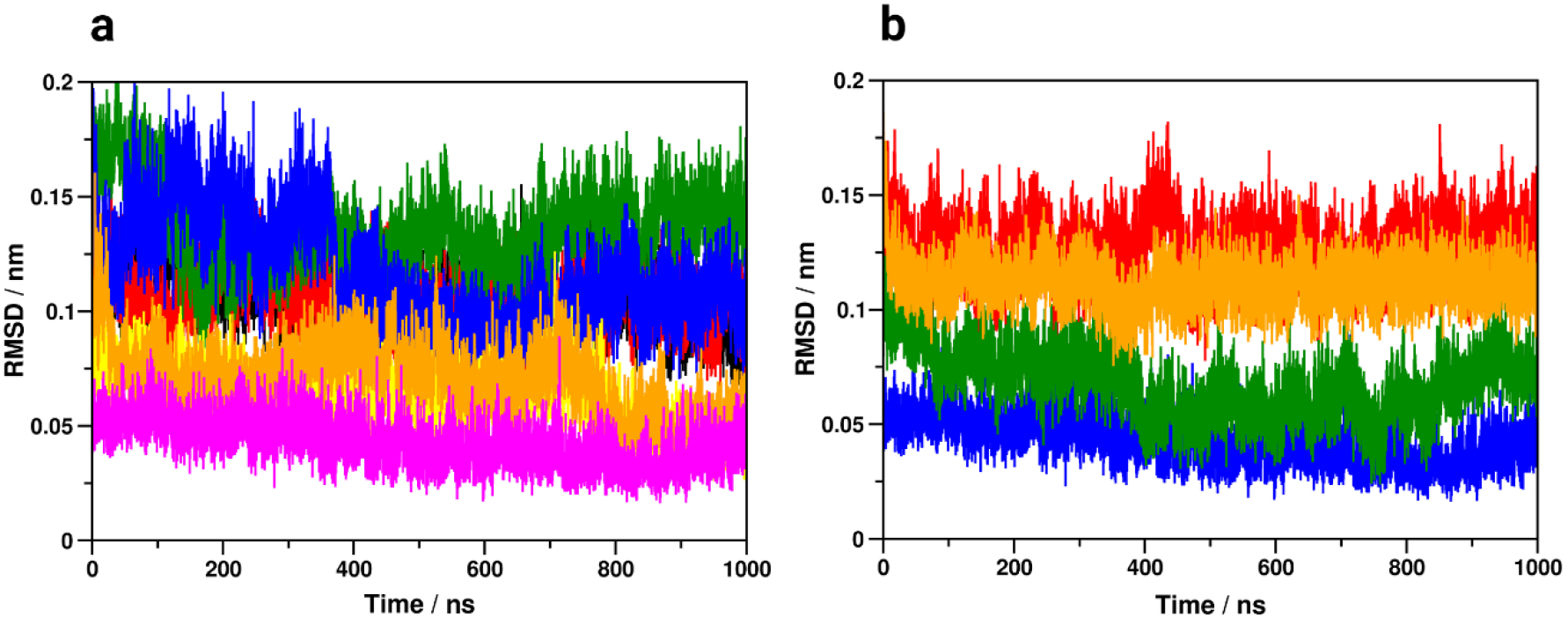
Properties of transmembrane helices. (a) Deviation from ideal α-helical geometry in the G_i_ protein-bound active state. Black: TM1, red: TM2, green: TM3, blue: TM4, yellow: TM5, orange: TM6, magenta: TM7. (b) Deviation of TM7 from ideal α-helical geometry in the active, G_i_ protein-bound (blue), beta-arrestin-2 bound (red), Nb39 nanobody-bound (green) and T4-lysozyme-fused (orange) states.

## CONCLUSION

The above results and considerations, as well as comparison to published mutation data have led us to suggest that large scale structural rearrangements may not be the key event of receptor activation. According to our alternative hypothesis, structural changes induced by agonist binding are subtle and the signal transduction mechanism is rather initiated by the perturbation of electrostatic balance within the binding pocket. Such perturbation is then propagated to the intracellular G protein-binding interface through the minuscule rearrangement of polar amino acid side chains of highly conserved structural motifs, located along TM7, and this charge shift is assisted by the inherent dipole moment of that helical segment. This alternative perspective of the activation mechanism, corroborated by a number of earlier indications,^24–36^ may help to explain ligand induced effects in multiple functional states. More importantly, this may highlight certain physico-chemical properties of ligands with different functional properties and may provide a new perspective for medicinal chemists in the pursuit of a new generation of GPCR drugs.

## ASSOCIATED CONTENT

### Supporting Information

The Supporting Information is available free of charge on the ACS Publications website at DOI: xxxxxxxxxx.

TM6 disposition analysis data. Secondary structure analysis data. Dynamic cross-correlation matrix analysis data. Polar signaling channel diagrams for all studied systems. Analysis results for intermolecular H-bonds and salt bridges. Side chain rotameric states analysis data for the polar signaling channel.

The following files are available free of charge.

Supporing Information (PDF)

## Author Contributions

A. M. built simulation systems, analyzed MD trajectories, executed multiple sequence alignment, prepared figures and wrote the manuscript. A. S. built simulation systems, analyzed MD trajectories and wrote the manuscript. M. R. Sz. performed glycosylation site prediction, built simulation systems and wrote the manuscript. A. B. conceptualized and supervised the study, performed molecular docking and MD simulations, analyzed MD trajectories, prepared images and wrote the manuscript with contributions from all other coauthors.

## ACKNOWLEDGMENT

Computing resources were provided by the National Institute of Informatics and Infrastructure, Hungary. The authors would like to thank Balázs Papp and Zoltán Lipinszki for their valuable suggestions with regard to the manuscript. M. R. Sz. was supported by the ÚNKP-19-3-SZTE-269 New National Excellence Program of the Ministry for Innovation and Technology.

## ABBREVIATIONS

GPCR: G protein-coupled receptor
TM6: 6^th^ transmembrane helix
ICL1: 1^st^ intracellular loop
H8: cytosolic helix
MD: molecular dynamics
EM2: endomorphin-2
TM: transmembrane
GTP: guanosine-triphosphate
ICL3: 3^rd^ intracellular loop
NVT: constant number of particles, volume and temperature
NPT: constant number of particles, pressure and temperature
RMSD: root mean square deviation
DCCM: dynamic cross-correlation matrix
MI: mutual information
ICL2: 2^nd^ intracellular loop
TM7: 7^th^ transmembrane helix
V_m_: transmembrane electrostatic potential

## REFERENCES

1. A. S. Hauser, M. M. Attwood, M. Rask-Andersen, H. B. Schiöth, D. E. Gloriam, Trends in GPCR Drug Discovery: New Agents, Targets and Indications. Nat. Rev. Drug Discov. 16, 829–842 (2017).

2. C. Munk, E. Mutt, V. Isberg, L. F. Nikolajsen, J. M. Bibbe, T. Flock, M. A. Hanson, R. C. Stevens, X. Deupi, D. E. Gloriam, An Online Resource for GPCR Structure Determination and Analysis. Nat. Methods 16, 151–162(2019).

3. N. R. Latorraca, A. J. Venkatakrishnan, R. O. Dror, GPCR Dynamics: Structures in Motion. Chem. Rev. 117, 139–155 (2017).

4. A. Kapoor, G. Martinez-Rosell, D. Provasi, G. de Fabritiis, M. Filizola, Dynamic and Kinetic Elements of m-Opioid Receptor Functional Selectivity. Sci. Rep. 7, 11255 (2017).

5. W. Huang, A. Manglik, A. J. Venkatakrishnan, T. Laeremans, E. N. Feinberg, A. L. Sanborn, H. E. Kato, K. E. Livingston, T. S. Thorsen, R. C. Kling, S. Granier, P. Gmeiner, S. M. Husbands, J. R. Traynor, W. I. Weis, J. Steyaert, R. O. Dror, B. K. Kobilka, Structural Insights into μ-Opioid Receptor Activation. Nature 524, 315–321(2015).

6. A. Koehl, H. Hu, S. Maeda, Y. Zhang, Q. Qu, J. M. Paggi, N. R. Latorraca, D. Hilger, R. Dawson, H. Matile, G. F. X. Schertler, S. Granier, W. Weis, R. O. Dror, A. Manglik, G. Skiniotis, B. K. Kobilka, Structure of the μ-Opioid Receptor-G_i_ Protein Complex. Nature 558, 547–552 (2018).

7. A. Manglik, A. C. Kruse, T. S. Kobilka, F. S. Thian, J. M. Mathiesen, R. K. Sunahara, L. Pardo, W. I. Weis, B. K: Kobilka, S. Granier, Crystal Structure of the m-Opioid Receptor Bound to a Morphinan Antagonist. Nature 485, 321–326(2012).

8. R. Sounier, C. Mas, J. Steyaert, T. Laeremans, A. Manglik, W. Huang, B. K. Kobilka, H. Déméné, S. Granier, Propagation of Conformational Changes During μ-Opioid Receptor Activation. Nature 524, 375–378(2015).

9. G. Fenalti, P. M. Giguere, V. Katritch, X. P. Huang, A. A. Thompson, V. Cherezov, B. L. Roth, R. C. Stevens, Molecular Control of μ-Opioid Receptor Signaling. Nature 506, 191–196(2014).

10. R. O. Dror, D. H. Arlow, P. Maragakis, T. J. Mildorf, A. C. Pan, H. Xu, D. W. Borhani, D. E. Shaw, Activation Mechanism of the b2-Adrenergic Receptor. Proc. Natl. Acad. Sci. USA. 108, 18684–18689(2011).

11. K. A. Marino, Y. Shang, M. Filizola, Insights into the Function of Opioid Receptors from Molecular Dynamics simulations of Available Crystal Structures. Br. J. Pharmacol. 175, 2834–2845 (2018).

12. R. A. Challiss, J. Wess, GPCR-G Protein Preassembly? Nat. Chem. Biol. 7, 657–658(2011).

13. M. J. Strohman, S. Maeda, D. Hilger, M. Masureel, Y. Du, B. K. Kobilka, Local Membrane Charge Regulates μ2-Adrenergic Receptor Coupling to Gi3. Nat. Commun. 10, 2234 (2019).

14. J. E. Zadina, L. Hackler, L. J. Ge, A. J. Kastin, A Potent and Selective Endogenous Agonist for the Mu-Opioid Receptor. Nature 386, 499–502(1997).

15. R. Gupta, E. Jung, S. Brunak, Prediction of N-Glycosylation Sites in Human Proteins. (2004), http://www.cbs.dtu.dk/services/NetNGlyc/.

16. P. Huang, C. Chen, W. Xu, S. I. Yoon, E. M. Unterwald, J. E. Pintar, Y. Wang, P. L. Chong, L. Y. Liu-Chen, Brain Region-Specific N-Glycosylation and Lipid Rafts Association of the Rat Mu Opioid Receptor. Biochem. Biophys. Res. Commun. 365, 82–88(2008).

17. A. Mann, S. Illing, E. Miess, S. Schulz, Different Mechanisms of Homologous and Heterologous μ-Opioid Receptor Phosphorylation. Br. J. Pharmacol. 172, 311–316(2015).

18. H. Zheng, E. Pearsall, D. Hurst, Y. Zhang, J. Chu, Y. Zhou, P. Reggio, H. Loh, P. Law, Palmitoylation and Membrane Cholesterol Stabilize μ-Opioid Receptor Homodimerization and G Protein Coupling. BMC Cell Biol. 13, 1–18(2012).

19. L. J. Pike, X. Han, K. N. Chung, R. W. Gross, Lipid Rafts are Enriched in Arachidonic Acid and Plasmenylethanolamine and Their Composition is Independent of Caveolin-1 Expression: A Quantitative Electrospray Ionization/Mass Spectrometric Analysis. Biochemistry 41, 2075–2088(2002).

20. H. I. Ingólfsson, M. N. Melo, F. J. van Eerden, C. Arnarez, C. A. Lopez, T. A. Wassenaar, X. Periole, A. H. de Vries, D. P. Tieleman, S. J. Marrink, Lipid Organization of the Plasma Membrane. J. Am. Chem. Soc. 136, 14554–14559 (2014).

21. W. Kabsch, C. Sander, Dictionary of Protein Secondary Structure: Pattern Recognition of Hydrogen-Bonded and Geometrical Features. Biopolymers 22, 2577–2637 (1983).

22. O. Lange, H. Grubmueller, Generalized Correlation for Biomolecular Dynamics. Proteins 62, 1053–1061(2006).

23. F. Sievers, A. Wilm, D. Dineen, T. J. Gibson, K. Karplus, W. Li, R. Lopez, H. McWilliam, M. Remmert, J. Söding, J. D. Thompson, D. G. Higgins, Fast, Scalable Generation of High-Quality Protein Multiple Sequence Alignments Using Clustal Omega. Mol. Syst. Biol. 7, 539

24. A. Jongejan, M. Bruysters, J. A. Ballesteros, E. Haaksma, R. A. Bakker, L. Pardo, R. Leurs, Linking Agonist Binding to Histamine H1 Receptor Activation. Nat. Chem. Biol. 1, 98–103(2005).

25. R. Liu, D. Nahon, B. le Roy, E. B. Lenselink, A. P. Ijzerman, Scanning Mutagenesis in a Yeast System Delineates the Role of the NPxxY(x)5,6F Motif and Helix 8 of the Adenosine A2B Receptor in G Protein Coupling. Biochem. Pharmacol. 95, 290–300(2015).

26. J. D. Hothersall, R. Torella, S. Humphreys, M. Hooley, A. Brown, G. McMurray, S. A. Nickolls, Residues W320 and Y328 within the Binding Site of the μ-Opioid Receptor Influence Opiate Ligand Bias. Neuropharmacology, 118, 46–58(2017).

27. S. C. Sealfon, L. Chi, B. J. Ebersole, V. Rodic, D. Zhang, J. A. Ballesteros, H. Weinstein, Related Contribution of Specific Helix 2 and 7 Residues to Conformational Activation of the Serotonin 5-HT2A Receptor. J. Biol. Chem. 270, 16683–16688(1995).

28. W. Xu, F. Ozdener, J. G. Li, C. Chen, J. K. de Riel, H. Weinstein, L. Y. Liu-Chen, Functional Role of the Spatial Proximity of Asp1142.50 in TMH 2 and Asn3327.49 in TMH 7 of the Mu Opioid Receptor. FEBS Lett. 447, 318–324(1999).

29. C. Galés, A. Kowalski-Chauvel, M. N. Dufour, C. Seva, L. Moroder, L. Pradayrol, N. Vaysse, D. Fourmy, S. Silvente-Poirot, Mutation of Asn-391 Within the Conserved NPxxY Motif of the Cholecystokinin B Receptor Abolishes Gq Protein Activation Without Affecting Its Association With the Receptor. J. Biol. Chem. 275, 17321–17327(2000).

30. L. S. Barak, M. Tiberi, N. J. Freedman, M. M. Kwatra, R. J. Lefkowitz, M. G. Caron, A Highly Conserved Tyrosine Residue in G Protein-Coupled Receptors is Required for Agonist-Mediated Beta 2-Adrenergic Receptor Sequestration. J. Biol. Chem. 269, 2790–2795(1994).

31. C. Prioleau, I. Visiers, B. J. Ebersole, H. Weinstein, S. C. Sealfon, Conserved Helix 7 Tyrosine Acts as a Multistate Conformational Switch in the 5-HT2C Receptor. Identification of a Novel “LOCKED-ON” Phenotype and Double Revertant Mutations. J. Biol. Chem. 277, 36577–36584 (2002).

32. I. Kalatskaya, S. Schüssler, A. Blaukat, W. Müller-Esterl, M. Jochum, D. Proud, A. Faussner, Mutation of Tyrosine in the Conserved NPxxY Sequence Leads to Constitutive Phosphorylation and Internalization, but Not Signaling of the Human B2 Bradykinin Receptor. J. Biol. Chem. 279, 31268–31276(2004).

33. W. G. Hol, Effects of the μ-Helix Dipole Upon the Functioning and Structure of Proteins and Peptides. Adv. Biophys. 19, 133–165(1985).

34. D. Bartuzi, A. A. Kaczor, D. Matosiuk, Interplay between Two Allosteric Sites and Their Influence on Agonist Binding in Human m Opioid Receptor. J. Chem Inf. Model. 56, 563–570(2016).

35. C. B. Pert, G. Pasternak, S. H. Snyder, Opiate agonists and antagonists discriminated by receptor binding in brain. Science 182, 1359–1361(1973).

36. O. N. Vickery, J. P. Machtens, G. Tamburrino, D. Seeliger, U. Zachariae, Structural Mechanisms of Voltage Sensing in G Protein-Coupled Receptors. Structure 24, 997–1007(2016).

37. M. P. Mahaut-Smith, J. Martinez-Pinna, I. S. Gurung, A role for membrane potential in regulating GPCRs? Trends Pharmacol. Sci. 29, 421–429(2008).

