## Supporting Information for "Correlated motions of conserved polar motifs lay out a plausible mechanism of G protein-coupled receptor activation"

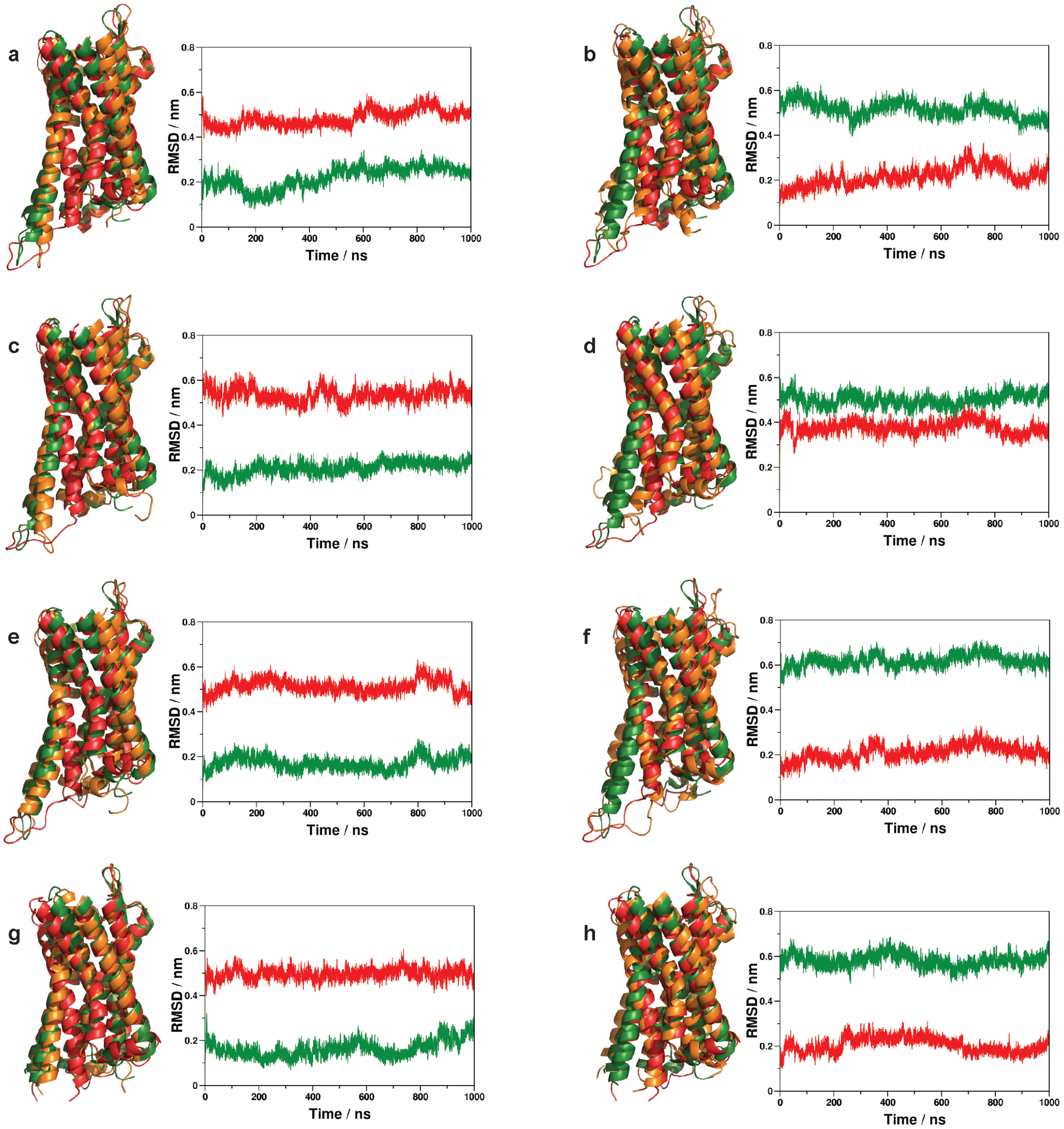


**Figure S1.** Disposition of TM6 during simulations with respect to the active (green) and inactive (red) crystallographic structures of the mu-opioid receptor. (a) Active receptor – $G_i$ protein complex. (b) Inactive receptor – $G_i$ protein complex. (c) Active receptor – beta-arrestin-2 complex. (d) Inactive receptor – beta-arrestin-2 complex. (e) Active receptor – Nb39 nanobody complex. (f) Inactive receptor – Nb39 nanobody complex. (g) Active receptor – T4-lysozyme fusion. (h) Inactive receptor – T4-lysozyme fusion.

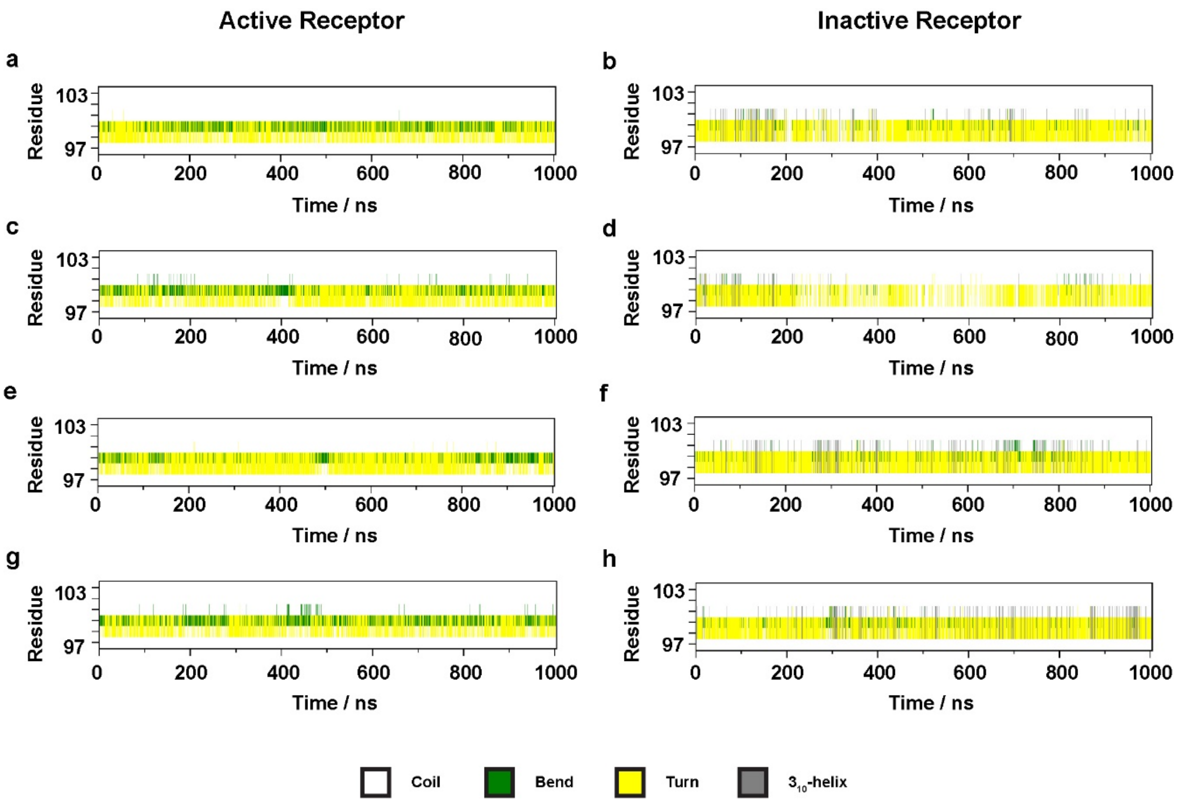


**Figure S2.** Evolution of the secondary structure of ICL1 during simulations. (a) Active receptor – $G_i$ protein complex. (b) Inactive receptor – $G_i$ protein complex. (c) Active receptor – beta-arrestin-2 complex. (d) Inactive receptor – beta-arrestin-2 complex. (e) Active receptor – Nb39 nanobody complex. (f) Inactive receptor – Nb39 nanobody complex. (g) Active receptor – T4-lysozyme fusion. (h) Inactive receptor – T4-lysozyme fusion.

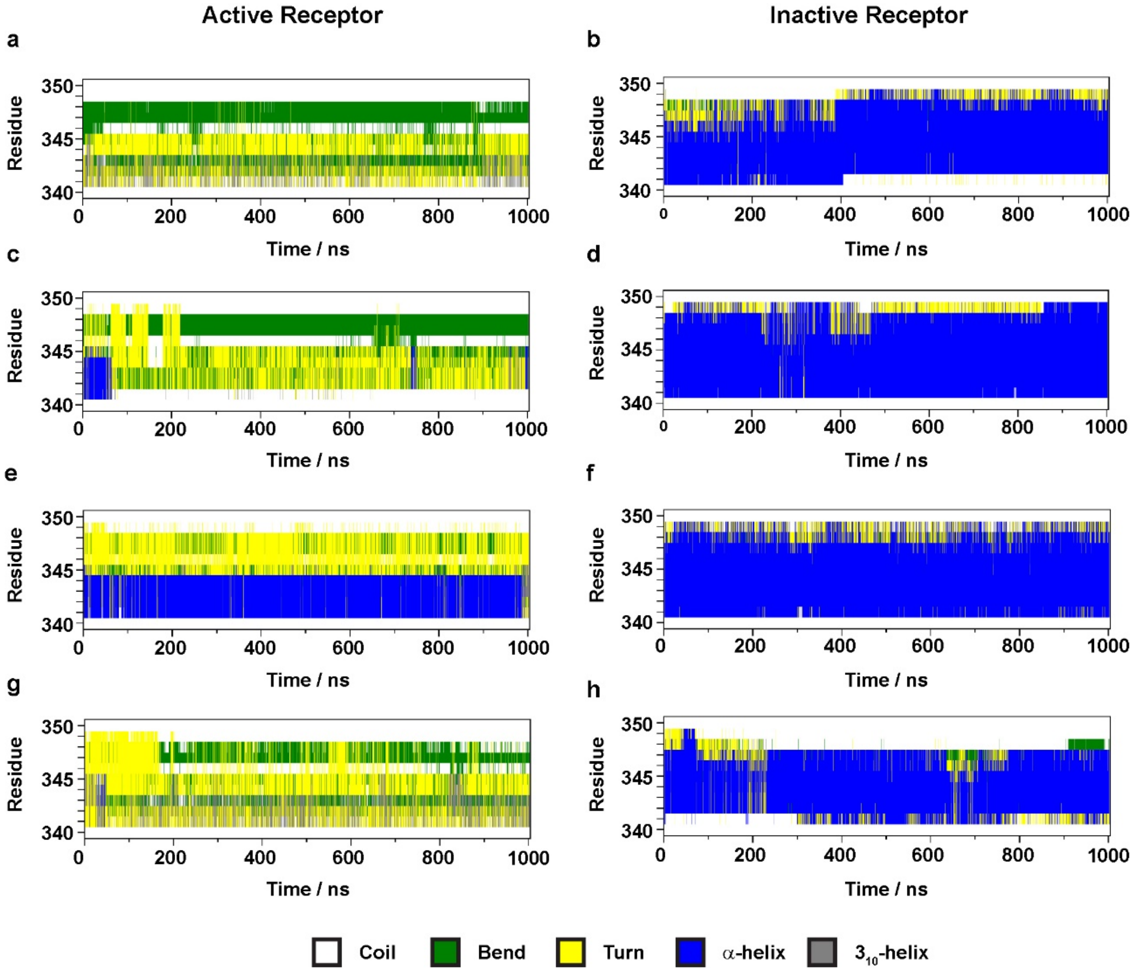


**Figure S3.** Evolution of the secondary structure of H8 during simulations. (a) Active receptor – $G_i$ protein complex. (b) Inactive receptor – $G_i$ protein complex. (c) Active receptor – beta-arrestin-2 complex. (d) Inactive receptor – beta-arrestin-2 complex. (e) Active receptor – Nb39 nanobody complex. (f) Inactive receptor – Nb39 nanobody complex. (g) Active receptor – T4-lysozyme fusion. (h) Inactive receptor – T4-lysozyme fusion.

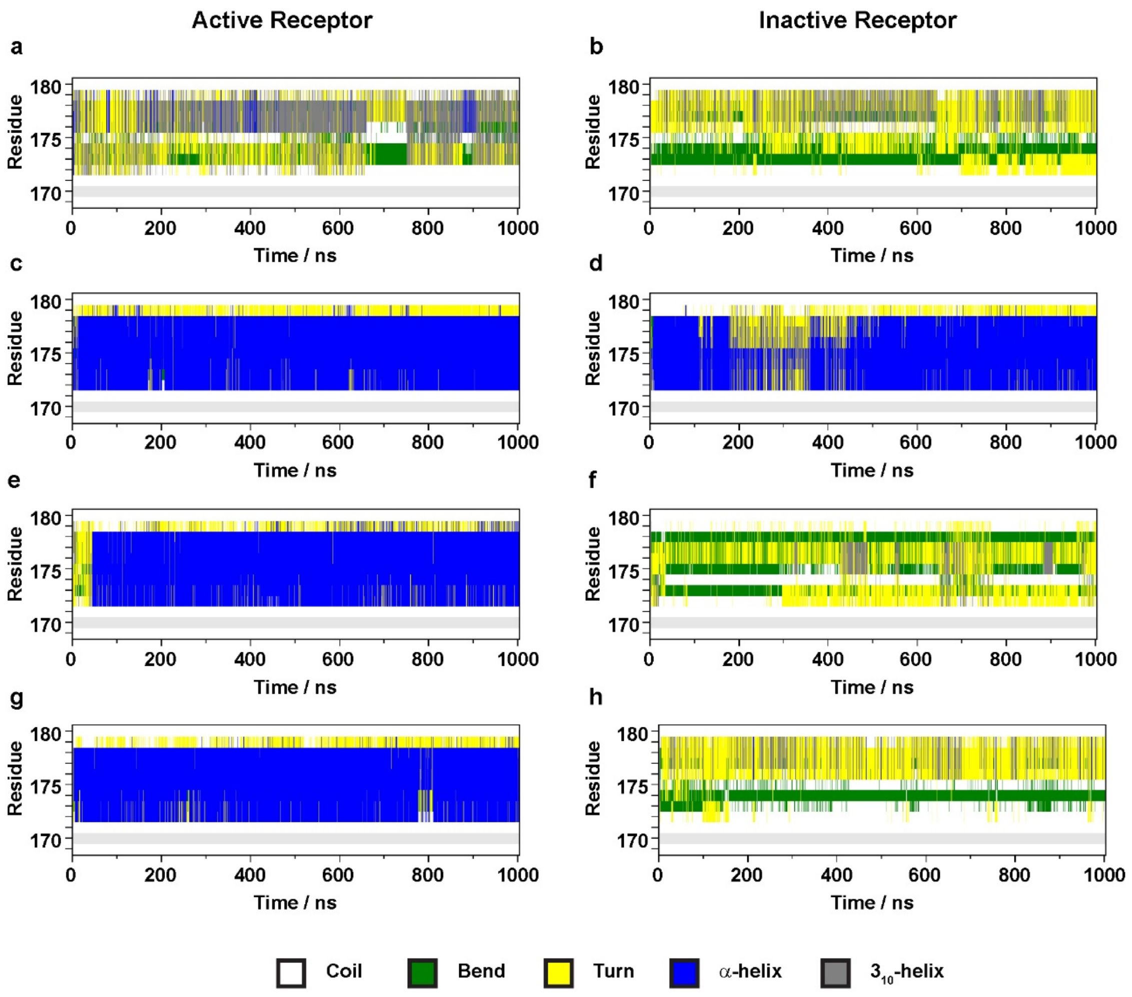


**Figure S4.** Evolution of the secondary structure of ICL2 during simulations. (a) Active receptor – $G_i$ protein complex. (b) Inactive receptor – $G_i$ protein complex. (c) Active receptor – beta-arrestin-2 complex. (d) Inactive receptor – beta-arrestin-2 complex. (e) Active receptor – Nb39 nanobody complex. (f) Inactive receptor – Nb39 nanobody complex. (g) Active receptor – T4-lysozyme fusion. (h) Inactive receptor – T4-lysozyme fusion.

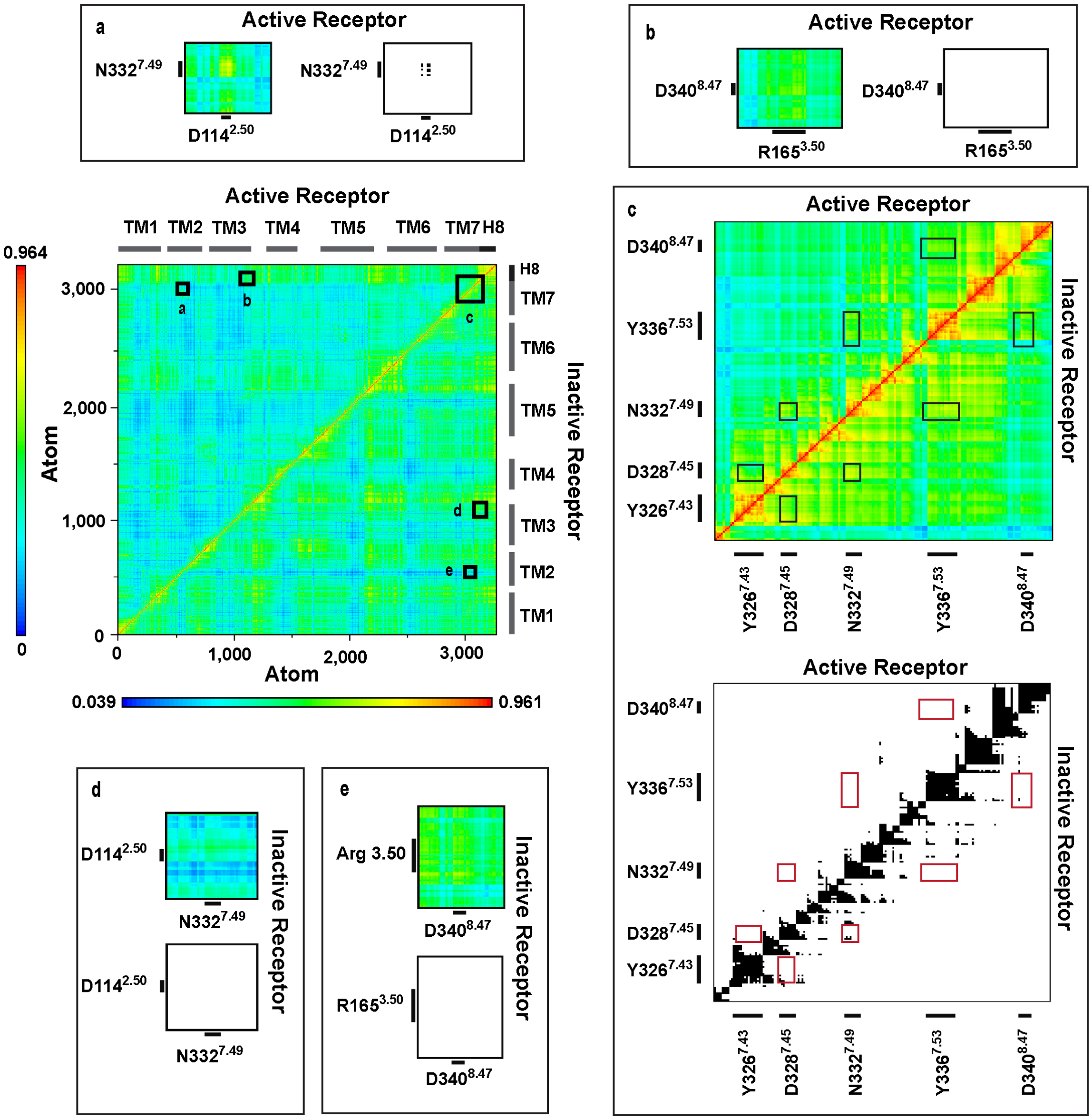


**Figure S5.** Dynamic cross-correlation matrices of the beta-arrestin-2-bound mu-opioid receptor in active and inactive states. Panels (a-e) are magnified views of regions of amino acid residues of interest. Black and white panels show correlations above the threshold of 0.7 MI.

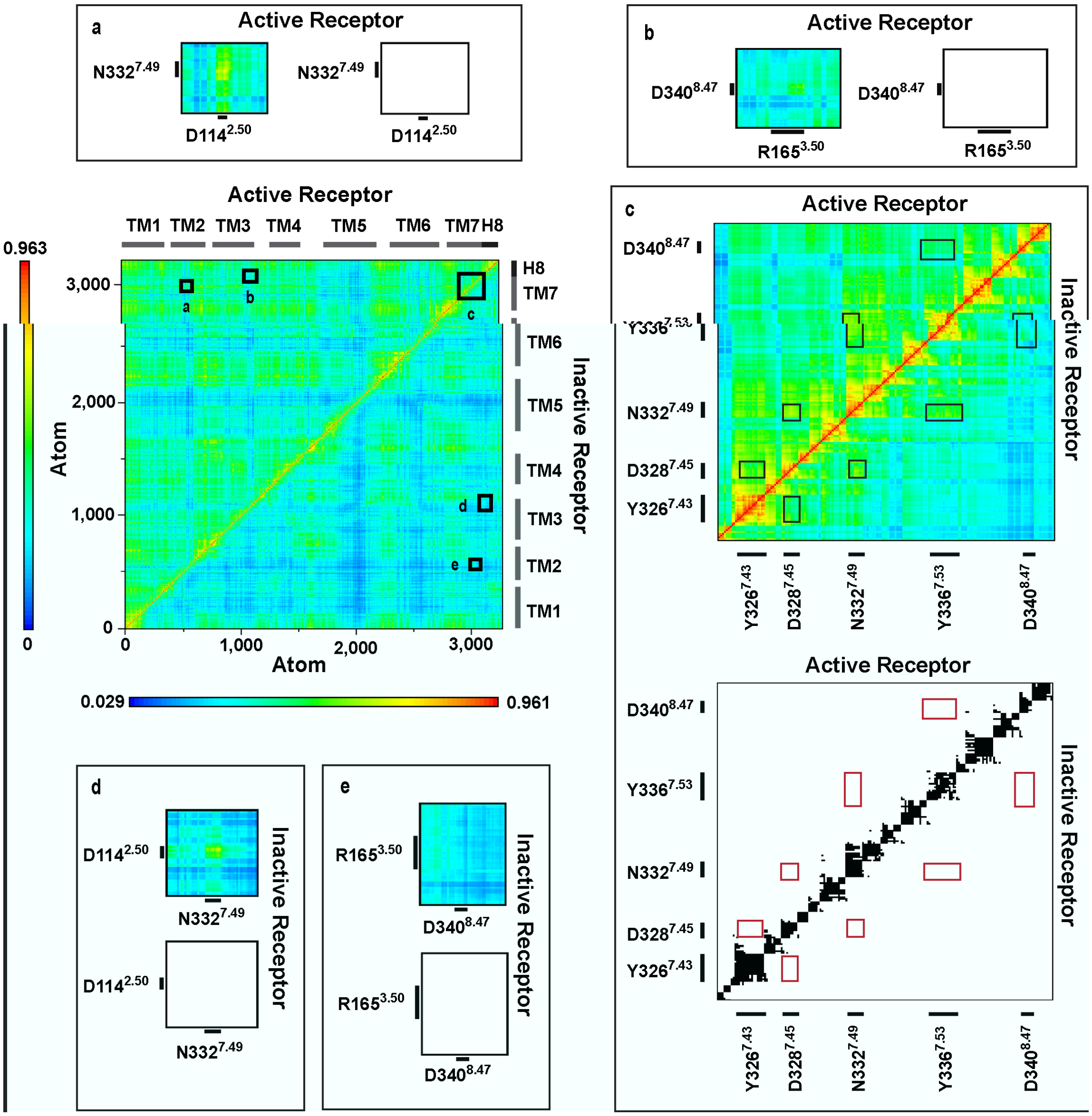


**Figure S6.** Dynamic cross-correlation matrices of the Nb39 nanobody-bound mu-opioid receptor in active and inactive states. Panels (a-e) are magnified views of regions of amino acid residues of interest. Black and white panels show correlations above the threshold of 0.7 MI.

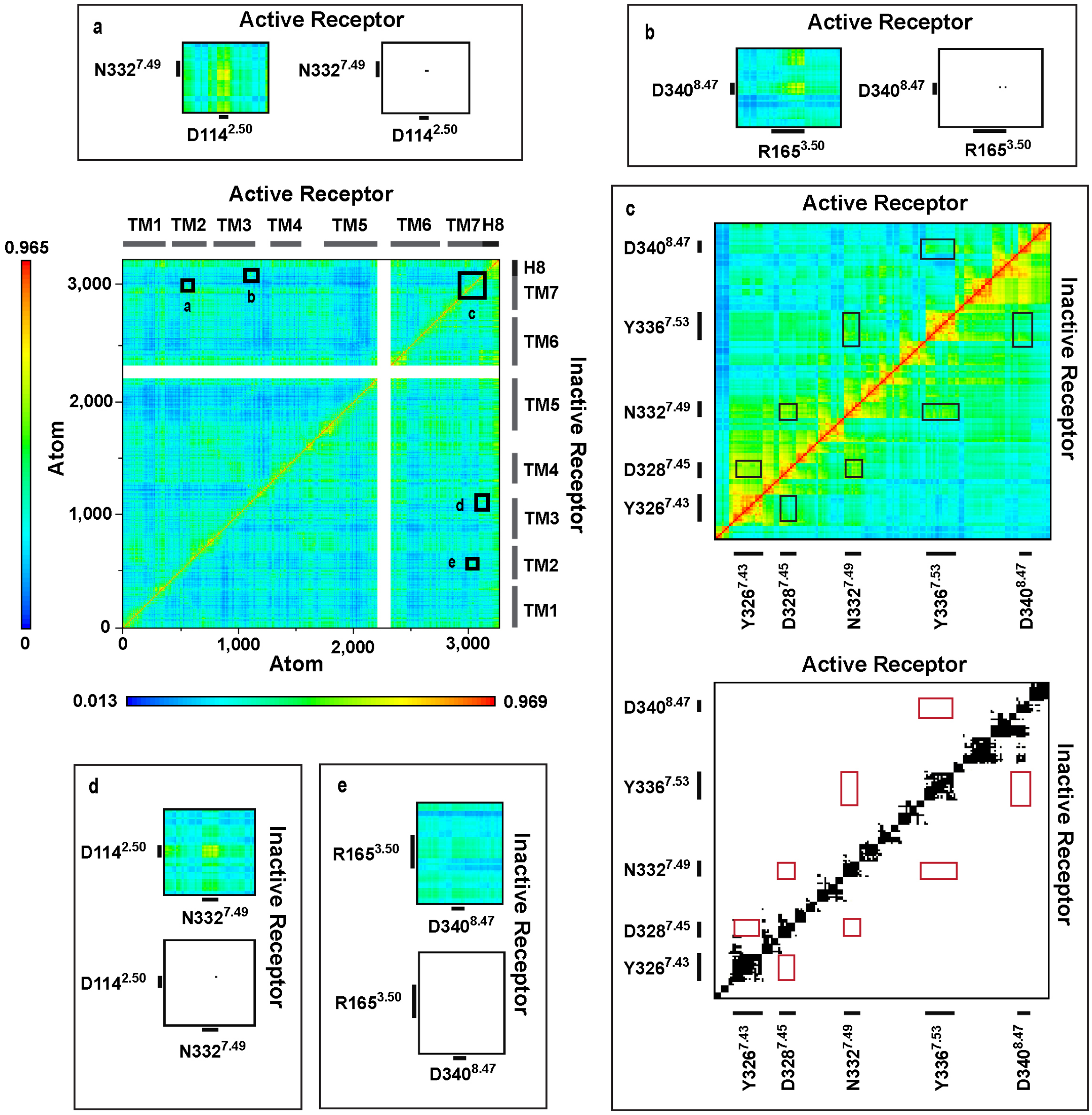


**Figure S7.** Dynamic cross-correlation matrices of the active and inactive mu-opioid receptor-T4-lysozyme fusion proteins. Panels (a-e) are magnified views of regions of amino acid residues of interest. Black and white panels show correlations above the threshold of 0.7 MI.

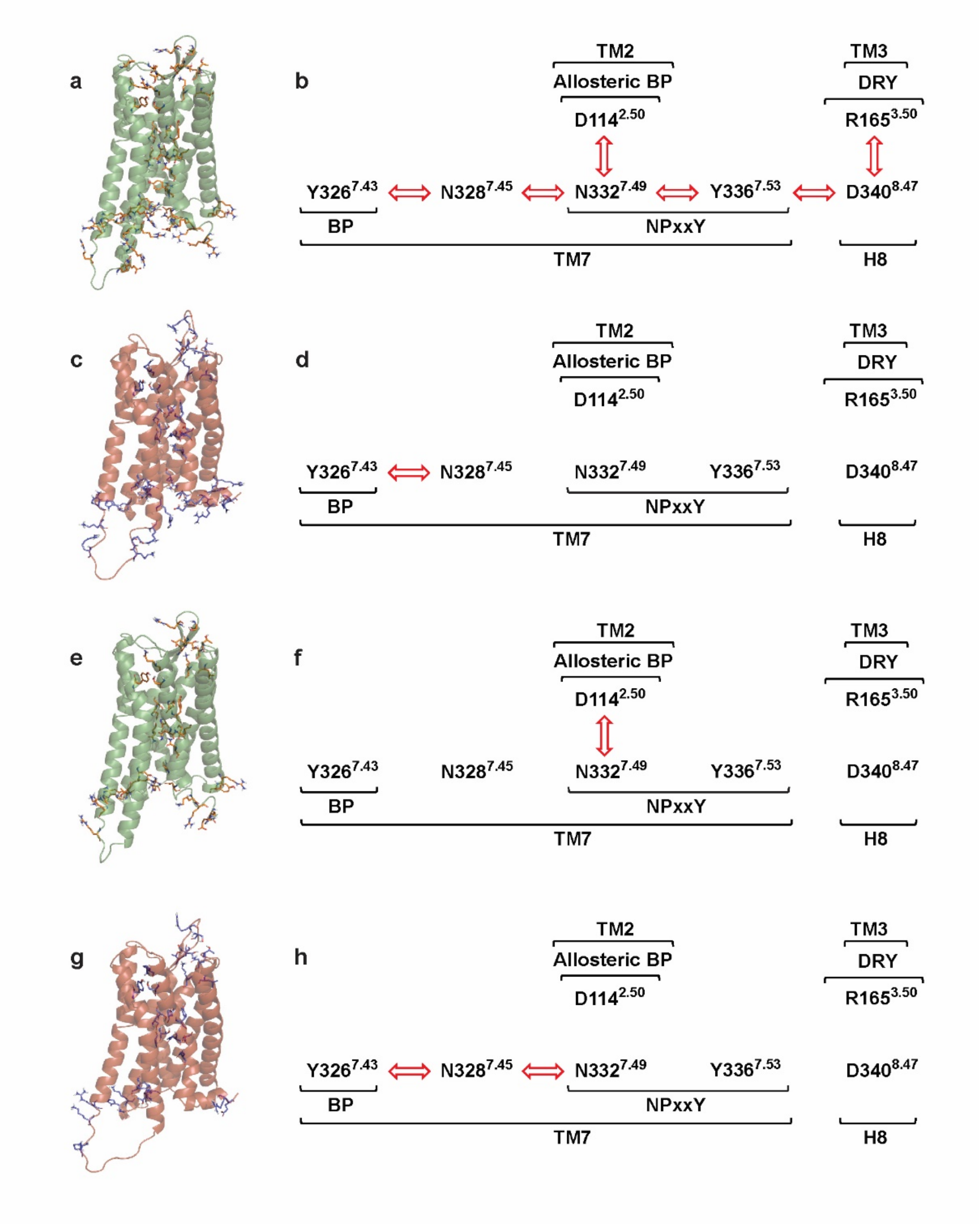


**Figure S8.** The polar signaling channel of the $G_i$ protein or beta-arrestin-2-bound mu-opioid receptor revealed by dynamic cross correlation analysis. Polar amino acids of which motions are correlated are shown in stick representation for (a) the $G_i$ protein-bound active, (c) $G_i$ protein-bound inactive, (e) beta-arrestin-2-bound active and (g) beta-arrestin-2-bound inactive states. Diagrams of channel residues in the (b) active receptor – $G_i$ protein, (d) inactive receptor – $G_i$ protein, (f) active receptor – beta-arrestin-2 and (h) inactive receptor – beta-arrestin-2 complexes. Red arrows indicate correlated motions of the respective amino acids.

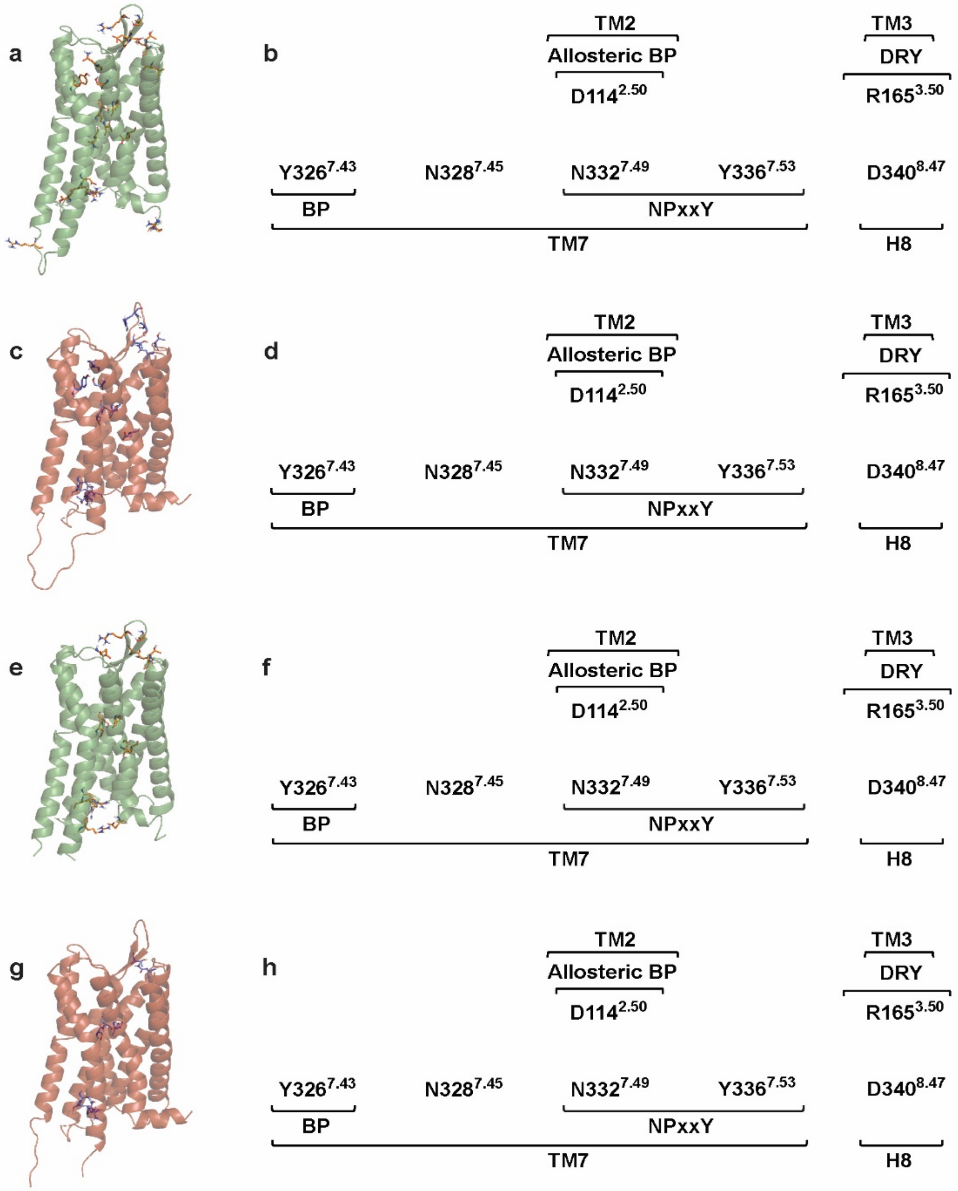


**Figure S9.** The polar signaling channel of the Nb39 nanobody-bound or T4 lysozyme-fused mu-opioid receptor revealed by dynamic cross correlation analysis. Polar amino acids of which motions are correlated are shown in stick representation for the (a) Nb39 nanobody-bound active, (c) Nb39 nanobody-bound inactive, (e) T4-lysozyme-fused active and (g) T4-lysozyme-fused inactive states. Diagrams of channel residues in the (b) active receptor – Nb39 nanobody, (d) inactive receptor – Nb39 nanobody, (f) active receptor – T4-lysozyme and (h) inactive receptor – T4-lysozyme complexes and fusions. Red arrows indicate correlated motions of the respective amino acids.

**Table S1.** Frequency of intermolecular salt bridges and H-bonds expressed as percentages of the total conformational ensemble, generated by MD simulations.

| Interactions | Residues involved, respectively | Active state | | | | Inactive state | | | |
|---|---|---|---|---|---|---|---|---|---|
| | | $G_i$ protein complex | Beta-arrestin-2 | Nb39 nanobody | Fused T4-lysozyme | $G_i$ protein complex | Beta-arrestin-2 | Nb39 nanobody | Fused T4-lysozyme |
| **Salt bridges** | | | | | | | | | |
| BP - EM2 | D147$^{3.32}$; Y1 | 68.4 | 76.2 | 60.0 | 59.2 | 11.7 | 3.6 | 7.0 | 1.3 |
| **H-bonds** | | | | | | | | | |
| BP - EM2 | D147$^{3.32}$; Y1 | 97.0 | 98.8 | 86.2 | 98.1 | 18.4 | 15.0 | 13.4 | 1.7 |
| ICL2 - H5 | V169-T180; K330-F354 | 23.0 | - | - | - | 3.8 | - | - | - |
| ICL2 - FL | V169-T180; G65-K78 | - | 40.8 | - | - | - | 1.4 | - | - |
| ICL2 - ML | V169-T180; P132-A140 | - | 86.4 | - | - | - | 87.3 | - | - |
| ICL2 - CL | V169-T180; L244-S246 | - | 19.3 | - | - | - | 0.0 | - | - |
| ICL2 - Nb39 | V169-T180; T52-V74 | - | - | 98.7 | - | - | - | 16.3 | - |

BP = orthosteric binding pocket of the mu-opioid receptor; EM2 = endomorphin-2; ICL2 = 2nd intracellular loop of the mu-opioid receptor; H5 = helix 5 of the $G_i$ protein [illegible] subunit; FL / ML / CL = finger loop / middle loop / C loop of beta-arrestin-2; Nb39 = Nb39 nanobody. Ballesteros-Weinstein numbering of residues is indicated in superscript.

**Table S2.** Side chain rotameric states and transitions of the residues of the polar signaling channel.

| | with the $G_i$ protein complex | | | | | | | | | | with beta-arrestin-2 | | | | | | | | | |
|---|---|---|---|---|---|---|---|---|---|---|---|---|---|---|---|---|---|---|---|---|
| | active | | | | | inactive | | | | | active | | | | | inactive | | | | |
| | rotamer | | | | | rotamer | | | | | rotamer | | | | | rotamer | | | | |
| residue | g+ | g- | t | $S^2$ | tr/ns* | g+ | g- | t | $S^2$ | tr/ns* | g+ | g- | t | $S^2$ | tr/ns* | g+ | g- | t | $S^2$ | tr/ns* |
| D114$^{2.50}$ | | 1.00 | | 0.98 | 0.00 | | 1.00 | | 0.98 | 0.05 | | 1.00 | | 0.99 | 0.00 | | 0.99 | 0.01 | 0.97 | 0.06 |
| R165$^{3.50}$ | | 0.98 | 0.02 | 0.95 | 0.05 | | 0.09 | 0.91 | 0.86 | 6.18 | | 1.00 | | 0.97 | 0.11 | | 0.98 | 0.02 | 0.93 | 1.04 |
| Y326$^{7.43}$ | | 1.00 | | 0.98 | 0.00 | | 1.00 | | 0.98 | 0.00 | | 1.00 | | 0.98 | 0.00 | | 1.00 | | 0.97 | 0.01 |
| N328$^{7.45}$ | | 0.06 | 0.94 | 0.84 | 0.61 | 0.11 | | 0.89 | 0.62 | 0.07 | | | 1.00 | 0.97 | 0.13 | | | 0.99 | 0.95 | 0.03 |
| N332$^{7.49}$ | | 1.00 | | 0.98 | 0.04 | | 1.00 | | 0.97 | 0.00 | | 1.00 | | 0.98 | 0.02 | | 0.14 | 0.86 | 0.67 | 0.12 |
| Y336$^{7.53}$ | | 1.00 | | 0.98 | 0.00 | | 1.00 | | 0.98 | 0.03 | | 1.00 | | 0.98 | 0.00 | | 1.00 | | 0.98 | 0.00 |
| D340$^{8.47}$ | | | 1.00 | 0.98 | 0.00 | | 1.00 | | 0.97 | 0.04 | | 0.34 | 0.66 | 0.48 | 0.26 | | 0.33 | 0.67 | 0.31 | 0.24 |
| I155$^{3.40}$ | 0.15 | 0.47 | 0.38 | 0.51 | 0.63 | | 0.15 | 0.85 | 0.79 | 0.20 | 0.97 | 0.03 | | 0.91 | 0.04 | 0.11 | 0.11 | 0.78 | 0.49 | 0.13 |
| F289$^{6.44}$ | 0.02 | 0.02 | 0.96 | 0.90 | 0.09 | | | 1.00 | 0.93 | 0.03 | | | 1.00 | 0.98 | 0.00 | | 0.96 | 0.04 | 0.98 | 0.01 |
| W293$^{6.48}$ | 0.01 | 0.50 | 0.49 | 0.31 | 0.01 | | 0.01 | 0.99 | 0.95 | 0.01 | 0.01 | 0.03 | 0.96 | 0.95 | 0.01 | 0.01 | 0.99 | | 0.98 | 0.01 |

| | with Nb39 nanobody | | | | | | | | | | with T4 lysozyme fusion | | | | | | | | | |
|---|---|---|---|---|---|---|---|---|---|---|---|---|---|---|---|---|---|---|---|---|
| | active | | | | | inactive | | | | | active | | | | | inactive | | | | |
| | rotamer | | | | | rotamer | | | | | rotamer | | | | | rotamer | | | | |
| residue | g+ | g- | t | $S^2$ | tr/ns* | g+ | g- | t | $S^2$ | tr/ns* | g+ | g- | t | $S^2$ | tr/ns* | g+ | g- | t | $S^2$ | tr/ns* |
| D114$^{2.50}$ | | 1.00 | | 0.98 | 0.00 | | 1.00 | | 0.98 | 0.01 | | 1.00 | | 0.98 | 0.00 | | 0.94 | 0.06 | 0.88 | 0.10 |
| R165$^{3.50}$ | | 1.00 | | 0.98 | 0.00 | | 1.00 | | 0.98 | 0.00 | | 1.00 | | 0.99 | 0.01 | | 1.00 | | 0.98 | 0.08 |
| Y326$^{7.43}$ | | 1.00 | | 0.98 | 0.00 | | 1.00 | | 0.98 | 0.00 | | 1.00 | | 0.98 | 0.01 | | 1.00 | | 0.98 | 0.00 |
| N328$^{7.45}$ | | 0.01 | 0.99 | 0.94 | 0.34 | | 0.18 | 0.82 | 0.57 | 0.22 | | | 1.00 | 0.96 | 0.09 | | 1.00 | | 0.87 | 0.21 |
| N332$^{7.49}$ | | 0.55 | 0.45 | 0.32 | 1.65 | | 0.99 | 0.01 | 0.96 | 0.03 | | 0.99 | 0.01 | 0.95 | 0.04 | | 0.03 | 0.97 | 0.92 | 0.01 |
| Y336$^{7.53}$ | | 0.99 | 0.01 | 0.96 | 0.15 | | 1.00 | | 0.98 | 0.00 | | 1.00 | | 0.98 | 0.01 | | 0.98 | 0.02 | 0.98 | 0.00 |
| D340$^{8.47}$ | | 0.10 | 0.90 | 0.75 | 0.07 | | 1.00 | | 0.97 | 0.01 | | | 1.00 | 0.98 | 0.01 | | 1.00 | | 0.96 | 0.00 |
| I155$^{3.40}$ | | 0.91 | 0.09 | 0.96 | 0.14 | | 0.05 | 0.95 | 0.92 | 0.15 | | 0.86 | 0.14 | 0.95 | 0.17 | 0.71 | 0.25 | 0.04 | 0.29 | 0.03 |
| F289$^{6.44}$ | | 0.01 | 0.99 | 0.96 | 0.00 | | 0.03 | 0.97 | 0.94 | 0.17 | | | 1.00 | 0.97 | 0.01 | | 0.88 | 0.12 | 0.97 | 0.02 |
| W293$^{6.48}$ | | 0.22 | 0.78 | 0.62 | 0.00 | 0.01 | 0.02 | 0.97 | 0.97 | 0.00 | | 1.00 | | 0.98 | 0.00 | | 1.00 | | 0.98 | 0.00 |

Data are shown only for populated rotameric states. Ballesteros-Weinstein numbering of residues is indicated in superscript.

* average number of transitions between rotameric states per nanosecond.
